# MAMMOTh: a new database for curated MAthematical Models of bioMOlecular sysTems

**DOI:** 10.1101/053652

**Authors:** Fedor Kazantsev, Ilya Akberdin, Sergey Lashin, Natalia Ree, Vladimir Timonov, Alexander Ratushny, Tamara Khlebodarova, Vitaly Likhoshvai

## Abstract

Living systems have a complex hierarchical organization that can be viewed as a set of dynamically interacting subsystems. Thus, to simulate the internal nature and dynamics of the whole biological system we should use the iterative way for a model reconstruction, which is a consistent composition and combination of its elementary subsystems. In accordance with this bottom-up approach, we have developed MAMMOTh (MAthematical Models of bioMOlecular sysTems) database that allows integrating manually curated mathematical models of biomolecular systems, which are fit to the experimental data. The database entries are organized as building blocks in a way that the model parts can be used in different combinations to describe systems with higher organizational level (metabolic pathways and/or transcription regulatory networks). The database supports export of single model or their combinations in SBML or Mathematica standards. The database currently contains more than 100 mathematical models for *Escherichia coli* elementary subsystems (enzymatic reactions and gene expression regulatory processes) that can be combined in at least 5100 complex/sophisticated models concerning such biological processes as: *de novo* nucleotide biosynthesis, aerobic/anaerobic respiration, and nitrate/nitrite utilization in *E. coli*. All current models are functionally interconnected and sufficiently complement public model resources.

**Database URL:** http://mammoth.biomodelsgroup.ru

## INTRODUCTION

Recent developments in experimental technologies have revolutionized biology and allowed researchers to generate huge amounts of highly resolved multiscale quantitative data in their respective biological fields. As a result, it caused a renaissance of the mathematical modeling of biological systems (1–4). Nowadays, mathematical modeling approaches are more often considered as general frameworks for the integration and analysis of experimental data and iterative investigation of dynamical biological systems (5–15). The general type of the model is determined on basis of the available information, used qualitative or quantitative data and the problem to solve. The assigned determinants for system behaviour define the required model variables. Mathematical models of dynamical biological systems can be formulated as systems of ordinary, partial, or delayed differential equations (ODE, PDE, DDE, respectively), stochastic equations, discrete operations or by hybrid models (4,13–23). The choice of the formalism usually depends on the complexity of biological system in question and availability of experimental data.

Currently, there are more than 1000 different published models of complex biomolecular systems (BMS), which are developed for several organisms (24–28). The number of such models is continuously increasing. However, reuse, expansion, or modification of the developed models is a labor-intensive process (29–32). Usually the models are designed as complete integrated systems and there are no tools available for researchers to modify their structural and/or functional contents in an automatic or semiautomatic way (31). The decomposition of the models into their elementary subsystems is an alternative approach to solve this problem. Such elementary subsystems can be viewed as “building blocks” and can be used to describe certain biomolecular functions and/or elementary processes of biological systems (14,33–36). The “functional network” represents a minimal model of the subsystem that is sufficient to perform a given function (37).

We have developed the MAMMOTh database which is designed to store mathematical models of an arbitrary biomolecular subsystem using the above-mentioned building blocks approach. The current version of the database contains more than 100 mathematical models of the *E.coli* biomolecular “elementary” subsystems, developed by authors of the present study using the original approach of generalized Hill’s functions (21). It should be mentioned that these “elementary” models then can be combined in various composite “complex” models, the total amount of which may be estimated at least to 5100 (which are consists of two and more connected “elementary” models). The models curated and stored in MAMMOTh are not presented in Biomodels.net (http://www.ebi.ac.uk/biomodels-main/) or Sabio-RK (http://sabio.villa-bosch.de) databases. MAMMOTh enables an automatic creation of mathematical models with different combinations of elementary subsystem. This database also allows export of the created models into different model standards to share them with other researchers and/or to integrate them in more complex models.

## RESULTS

### MAMMOTh database

The MAMMOTh database is designed for storage and accumulation of *elementary subsystem models* of biomolecular systems. *Elementary subsystem model* is the “functional network” model where the entities of the network and the mathematical model parameters are specified. The dynamics of each *elementary subsystem model* is related to published experimental data, which are referenced in the corresponding “model description” field. The database currently contains more than 100 mathematical models of BMS elementary subsystems: enzymatic reactions and gene expression regulation processes of *de novo* nucleotides biosynthesis, aerobic/anaerobic respiration and nitrate/nitrite utilization processes in *E.coli* cell. Every model consists of three parts: (1) the structural model (Figure 1a), which describes entities of the process, stoichiometric parameters and links on the public sources (EcoCyc, KEGG, PubChem); (2) the mathematical model (Figure 1b), which describes rate equation; (3) the kinetic parameter values (Figure 1c). Each mathematical model in the database has the description field where one can find references on articles which were the basis for the model development or which contain a quantitative experimental data used for model parameters fitting. The mathematical models were manually curated and verified by expert authors. The majority of the models were reconstructed using generalized Hill functions (21). These functions enable flexible formulation of model equations by solving structural and parametric inverse problems optimally aligning the model complexity with the level of detail in the relevant experimental data.

**Figure 1.**
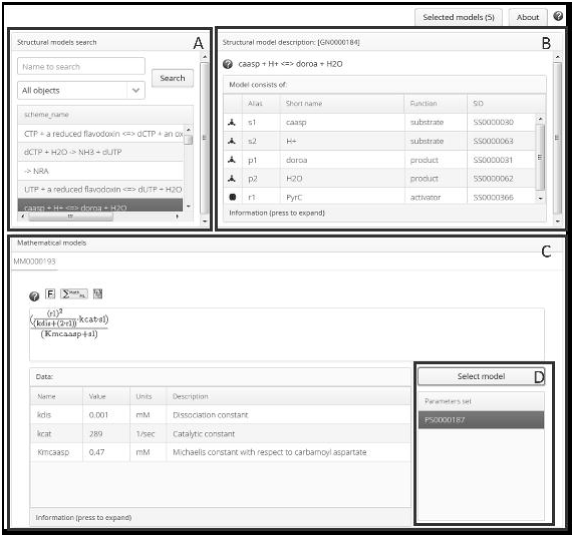
The MAMMOTh interface. (a) The query box. (b) The biochemical preview of the BMS elementary subsystem with the information about participants and model itself. (c) The mathematical model description of the selected subsystem. There are mathematical equation that you can download in different standards, model parameters and their values. (d) The area of available parameters value sets. The «select model» button is for moving of the selected model with certain parameters values into the «selected models» list. Each part has the «information» tab that contains additional information for selected entity as well as links to external sources.

To check the uniqueness of mathematical models in MAMMOTh database we have compared model variables (e.g., metabolites, enzymes) with model variables from other open access resources, such as "Sabio-RK” (38), “Biomodels.net” (39), CellML Model Repository (26) and “JWS online” (24). For this purpose we have used either APIs of the corresponding services or the manual data extraction (JWS online). PubMed IDs were used in the intersection search between articles that used for the model development or that contained experimental data for the model parameters fitting. We found two articles, which are also used in Sabio-RK to describe kinetic parameters (Pubmed: 1659321, 10074342; MAMMOTh links are [GN0000093; MM0000129] and [GN0000185; MM0000194], respectively).

To compare with data in *Biomodels.net* repository, we used the two-level procedure as follows. First, we used Pubmed ID for articles, which describe models or experimental data exploited for model adaptation. Then we compared database contents using Enzyme Commission numbers (EC number). At the first comparison level we have found three reference intersections only, which were articles with description of experimental data: 1) This case is PubMed ID: 19561621 (40), where several metabolites concentration and enzymatic activities data for the *E.coli* cell were used for the model of central metabolic pathway in the Biomodels.net (BIOMD0000000244) (41) under the investigation of bacterial cell adaptation to environment conditions. But the same data was used for another model in the MAMMOTh (GN0000141) to study the mechanism of the enzymatic reaction, which is catalyzed by NADH dependent nitrate reductase (EC:1.7.1.4); 2) This case is PubMed ID:9202467 (42) where *cAMP* concentration in dependence on glucose concentration was used for reconstruction and adaptation of central cell metabolic pathway model in Biomodels.net (BIOMD0000000051) (17) while the same data was used for exploration *guaA* gene expression regulation in the MAMMOTh (GN0000236); 3) the last case is PubMed ID:2200929 (43) which is the review article dedicated to metabolite and enzymes concentrations in cell of the *E. coli*, yeast, *Dictyostelium discoideum* and others. The data were used for model reconstruction of enzymatic reaction, catalyzed by phosphofructokinase in the pancreatic beta-cells in the Biomodels.net (BIOMD0000000225, BIOMD0000000236) (44). At the same time the quantitative data were used for exploration of N-carbamoyl-L-aspartate biosynthesis in the MAMMOTh model (GN0000183). Thus, the first step of the comparative analysis has shown that there are only three intersections between Biomodels.net and MaMoNT databases on experimental data, which were used for reconstruction of different biological processes.

The comparative analysis performed with the EC numbers at the second step gave us 11 general cases. The majority of the cases in Biomodels.net are associated with another organisms (mouse, human) (45), another cell type (46) or associated with the pipelines/techniques for the automation of model reconstruction process (47) (the experimental data were not mentioned in them).

Intersections in references on articles between MAMMOTh database and both JWS online and CellML systems were not found. Eventually, we have presented approximately 100 mathematical models of BMS elementary subsystems which were adapted to experimental data and were not published before in articles (besides of own reviewed abstracts in the conference proceedings) (Figure. 2).

**Figure 2.**
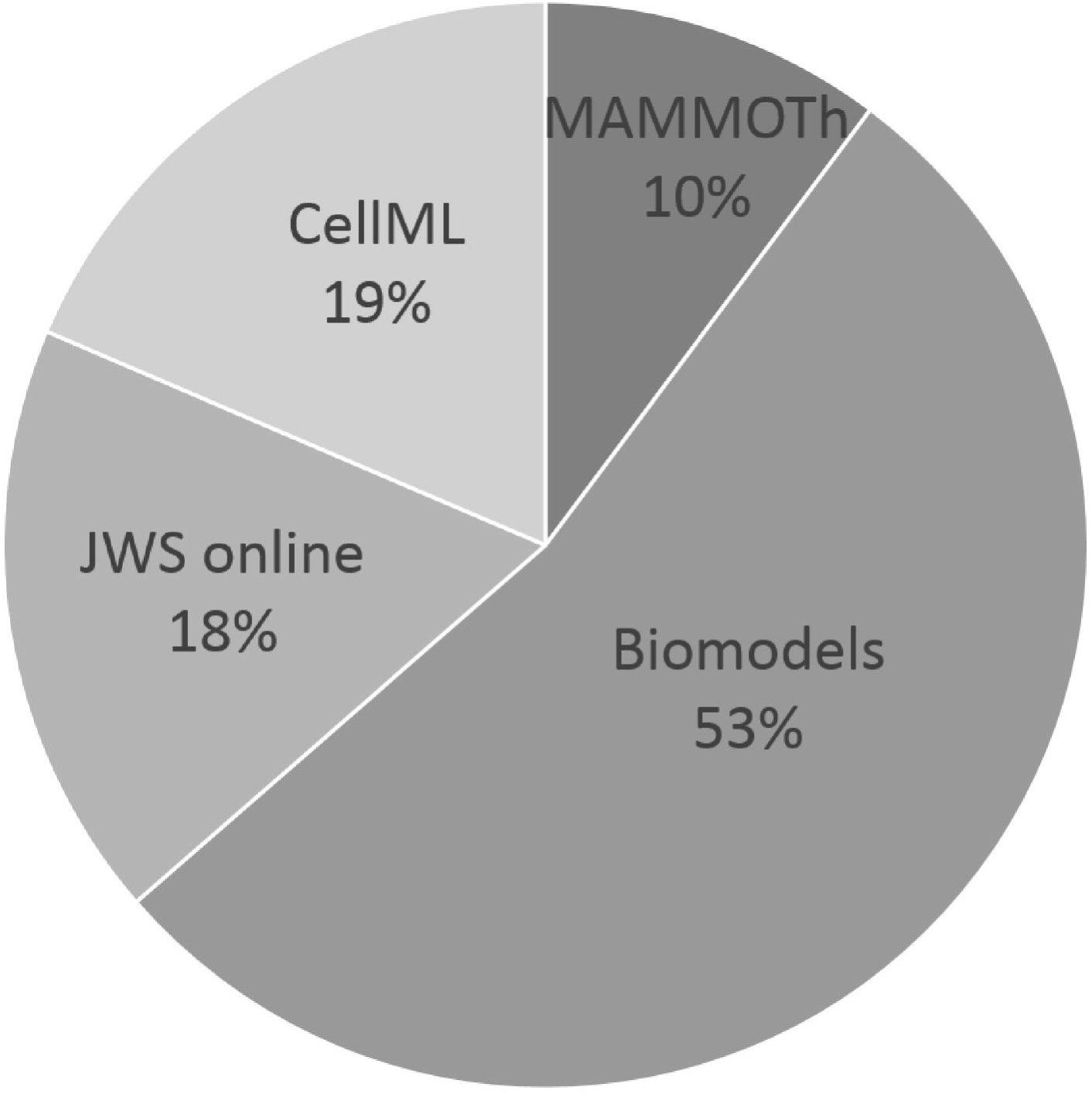
The quantity of mathematical models that are in open access and validated by experts.

### MAMMOTh software architecture

The MAMMOTh has been developed as Vaadin (vaadin.com) application with PostgreSQL database. It provides an access to the repository of elementary subsystems models by means of search for name of substances involved in the inquired subsystem, IDs of available kinetic models describing the process and IDs of the corresponding parameter sets. The database has been developed using the hybrid approach between relation and XML ones. The relational database provides fast access to models but the BMS subsystems are better to keep as XML, due to its benefits of the structural organization. Combination of approaches gave us the flexible database structure and simple data search requests (Figure 3).

**Figure 3.**
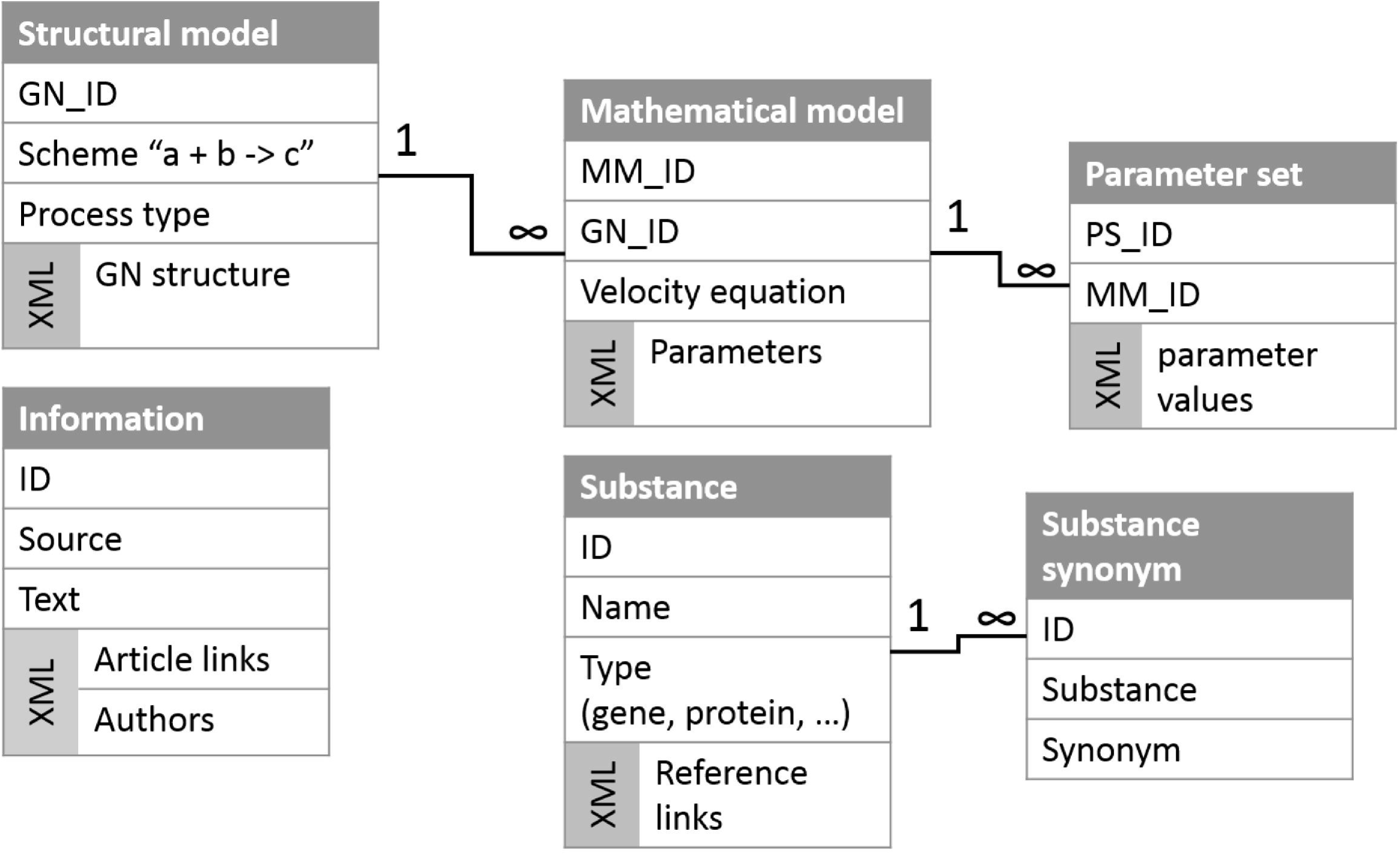
The MAMMOTh database scheme.

The data storage system enables us to utilize additional features. Each model in the database has been developed on basis of the general entities vocabulary. It makes possible to compare entities in the different models by identifiers directly. Using this property, the MAMMOTh provides some functionality for manipulation with models. To follow the “building blocks” strategy for complex models reconstruction we have implemented a method of template models generation reconstructed by the automatic composition from a subset of elementary models which are presented in the database. These assembled models can be exported in such well known formats as SBML, MATHEMATICA and Pajek (48) for further analysis in other tools.

#### Access and search routine

The web-interface of the MAMMOTh enables the user to search for models by names and types (gene, RNA, protein and substance) of entities (Figure 1a). As a result of the search the system generates a list of structural models, the components of which have a match in at least one of synonyms. When the user selects a structural model from the resulting list, a window opens containing lists of substrates, products, regulators of the elementary subsystems with relevant links to other database as well as a set of mathematical models. Rate reaction functions are visualized as mathematical formulas. Each model has an information field with the model description and links to relevant publications as well as a list of parameter sets in the section describing the mathematical model. A several sets of parameter values can be stored for the individual model in the database.

#### Models integration routine

The certain set of parameters can be selected by clicking the «select model» button which in turn allows the user to transfer mathematical model in the section «selected models» in upper right corner of the application window (Figure 1d). The user compiles a list of elementary models by the scheme described above. Additionally, the graph view of whole complex model is available too (Figure 4).

**Figure 4.**
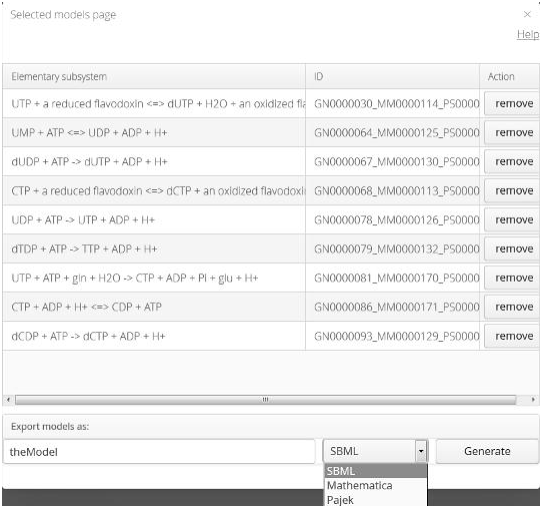
The interface for selected models. The “Generate” button activates the export of the complex model into the selected standard: SBML, Mathematica or Pajek.

#### Model export

After completion of the selection procedure the user can reconstruct a mathematical model from elementary models by selecting the appropriate format and clicking the «generate» button. Models are automatically assembled by the formation of global rates of component concentration (model variable) changes. The global rate represents a sum of the rates of the model variable changes in the selected elementary models. The MAMMOTh supports export of the BMS elementary subsystem models or models bunch, integrated into the complex one model, to the well-known standards: SBML, Mathematica and Pajek (48) (Figure 4). Due to the fact that each model in the database was adapted to the experimental data, further investigation will take less time to solve the parameters fitting (inverse) problem for the exported integrative model. The models bunch integrated into the one model could be a «growth point» in the process of complex models development and fitting or it can be just a part for more extended model(s) being developed in any other tool. We erase the support model information during the export to ensure non-conflicts with the selected standard.

## DISCUSSION

The hierarchical organization of biomolecular systems and elaborated “building blocks” approach enable systems biologists to iteratively develop integrative models by means of consistent composition of elementary models (Figure 5) (13,49–54). The elementary subsystems in the MAMMOTh database can be used in mathematical models of different biological systems.

**Figure 5.**
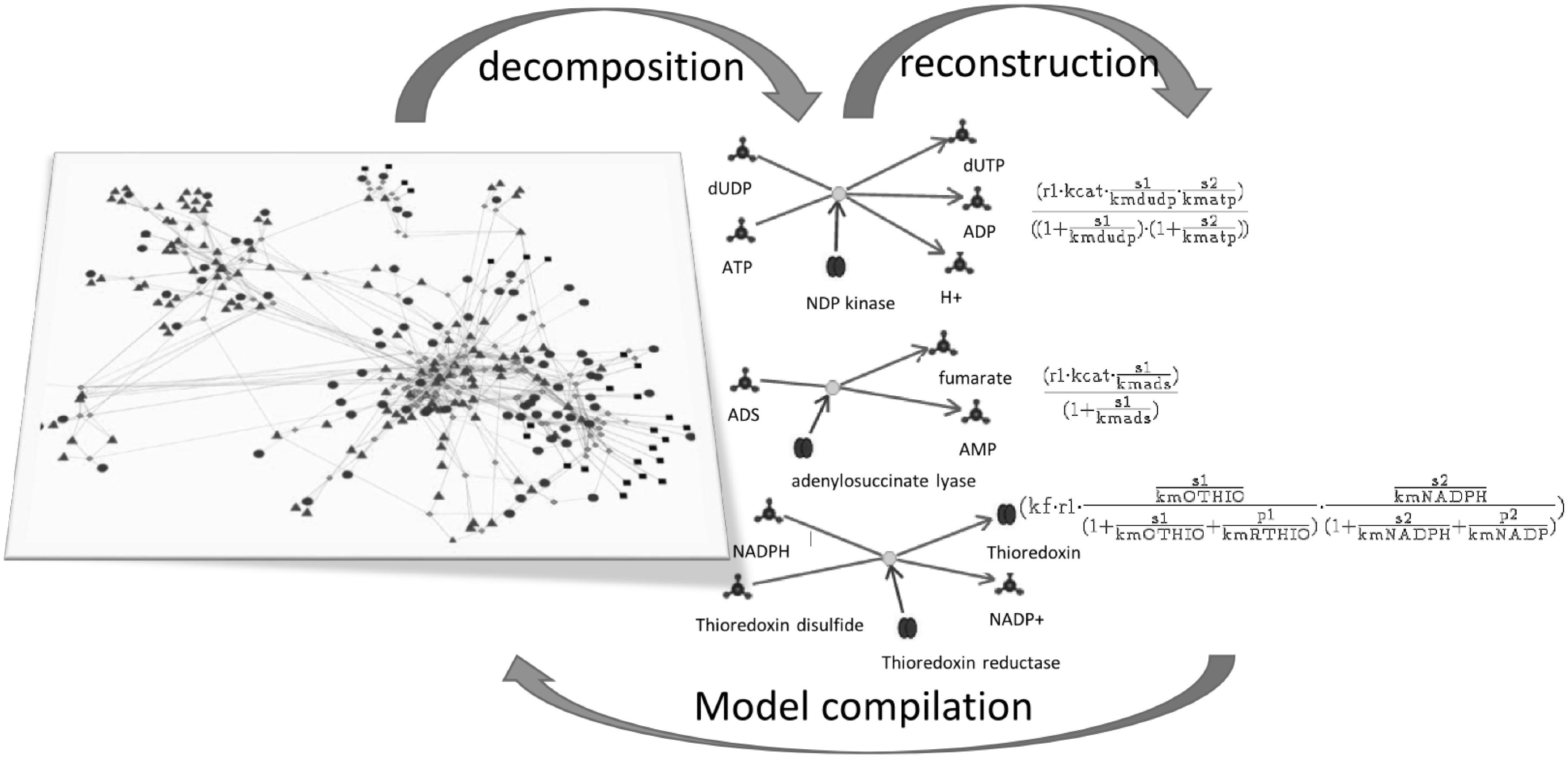
The scheme of model reconstruction process in terms of “building blocks”.

However, a purely mathematical approach can be more often found to the study of conserved regulatory circuits - "functional networks", which represent the minimal models of subsystems sufficient to perform a given function. These functional networks, aggregated in the database, (37) are extremely useful in the BMS model reconstruction process. These networks can be used to build preliminary “scratch” models, but the models obtained are rather mathematical abstractions. They describe interactions of biological entities, their regulatory effects that lead to the appropriate dynamics in the selected parameters constraints. But the model dynamics is sensitive to the variation of parameters values (30,55). Only the use of a physiologically appropriate parameters of the model allows researchers to reproduce the experimentally observed dynamics of the considered living system (30,55).

As an example of the *“building blocks”* approach, the most cited *Biomodels.net* database (56) was extended by means of 140 000 models of BMS subsystems models automatically generated by *Path2models* tool (57). The generation was based on structural and functional descriptions which were extracted from *KEGG* database (58). Meanwhile, less than 600 mathematical models in the *Biomodels.net* database are associated with papers and curated by experts. It means that together with the databases of “functional networks” abstract models (37,51,57,59–61) the databases of adapted models describing the dynamical patterns of biological subsystems on different hierarchical levels should be developed. In the future, it will allow us to replace the template models on fitted ones in the repositories and will give the opportunity for a full-scale reconstruction of models for different types of cells, tissues or even whole organisms. Karr and coauthors have already implemented this philosophy in the whole genome *Mycoplasma genitalium* model (4). It led to the development of the *WholeCellKB* (http://wholecellkb.stanford.edu) database which contains the description of functional cell subsystems.

## CONCLUSIONS

We have presented the MAMMOTh database of the kinetic models where the elementary models (enzymatic reaction or the gene expression regulatory process) and integrated complex models (metabolic pathways) can be exported in the well-known standards for model analysis: SBML, Mathematica and Pajek (48). Today the database contains more than 100 BMS elementary subsystems mathematical models of the enzymatic reactions describing *de novo* nucleotide biosynthesis, aerobic/anaerobic respiration and nitrate/nitrite utilization in *Escherichia coli*, which have not been presented in public BMS mathematical models resources.

Each model in the database is fitted to experimental data (data was taken from over 140 sources) and manually curated by experts in systems biology. The developed elementary models have been used for reconstruction of a several more complex BMS models (62–68).

The MAMMOTh repository has a friendly user interface providing the manual models query and the REST API that can be used for programmatic data access and the integration with external software tools. Currently, there are a few tools which give the opportunity to generate the template mathematical models on basis of the structural description of simulated biochemical systems systems (57,69–74). We strongly believe that the developed database carrying the fitted mathematical models of BMS elementary subsystems will impact on and improve the reliability of the model automatic generation process.

## AVAILABILITY

The MAMMOTh is freely available for the academic use: http://mammoth.biomodelsgroup.ru. REST API: http://mammoth.biomodelsgroup.ru/api

## LIST OF ABBREVIATIONS

DDE – Delay Differential Equation.

BMS – Biomolecular Systems.

MAMMOTh – MAthematical Models of bioMOlecular sysTems.

ODE – Ordinary Differential Equations.

PDE – Partial Differential Equations.

## AUTHORS’ CONTRIBUTIONS

FK has developed the MAMMOTh software including database and GUI levels. VT has developed the MAMMOTh GUI system. IA, NR, SL and AR have developed series of mathematical models, performed their analysis, checking and inserting it into the database. VL and TK performed analysis of the models, VL and AR supervised the development and implementation of the models. FK, IA, SL and AR wrote the manuscript with editing by TK and VL. All authors read and approved the final manuscript.

